# Emergence of a node-like population within an in vitro derived Neural Mesodermal Progenitors (NMPs) population

**DOI:** 10.1101/326371

**Authors:** Shlomit Edri, Penelope Hayward, Wajid Jawaid, Alfonso Martinez Arias

## Abstract

The mammalian embryos Caudal Lateral Epiblast (CLE) harbours bipotent progenitors, called Neural Mesodermal Progenitors (NMPs), that contribute to the spinal cord and the paraxial mesoderm throughout axial elongation. Here we performed a single cell analysis of different in vitro NMPs populations produced either from embryonic stem cells (ESCs) or epiblast stem cells (EpiSCs) and compared them to E8.25 CLE mouse embryos. In our analysis of this region our findings challenge the notion that NMPs should coexpress *Sox2* and *T*. We built a Support Vector Machine (SVM) based on the embryo CLE and use it as a classification model to analyse the in vitro NMP-like populations. We showed that ESCs derived NMPs are heterogeneous and contain few NMP-like cells, whereas EpiSCs derived NMPs, produce a high proportion of cells with the embryo NMP signature. Importantly, we found that the population from which the Epi-NMPs are derived in culture, contains a node-like population, which is responsible for maintaining the expression of *T* in vitro. These results mimic the events in vivo and suggest a sequence of events for the NMPs emergence.

## Introduction

In mammalian embryos, the trunk consists of the endoderm, the spinal cord and the derivatives of different kinds of mesodermal (axial, paraxial, intermediate and lateral plate). Much of our current understanding regarding the development of this body region has focused on two progenitor cell populations: the node, that will give rise to the axial mesoderm and the Neural Mesodermal Progenitors (NMPs), a bipotent stem cell population that contributes to the spinal cord and the paraxial mesoderm (PXM) (Henrique et al., 2015; Selleck and Stern, 1991; Wilson et al., 2009). Both populations are closely related within the anterior region of the Caudal Epiblast (CE) in the embryo (Wymeersch et al., 2016). This association persists for as long as the node is visible, between stages E7.5 and E9.0 (Fig. 1, Fig. S1 and (Wymeersch et al., 2016; Yamanaka et al., 2007). It is not clear when the NMPs arise but their association with the node suggests that they might emerge at the same time, around E7.5; the NMP population must then proliferate to sustain the axial extension process. Absence of the node results in severe axial truncations (Ang and Rossant, 1994; Davidson and Tam, 2000; Weinstein et al., 1994), suggesting a relationship between the node and the establishment and maintenance of the NMPs. However, little is known about these interactions.

The earliest identifiable NMPs emerge in the CE of E8.25 embryos distributed between the Node Streak Border (NSB) and the Caudal Lateral Epiblast (CLE) (Cambray and Wilson, 2007; Wymeersch et al., 2016). They are associated with the coexpression of *T* (*Brachyury)*, *Sox2* and *NKx1-2* (Henrique et al., 2015; Steventon and Martinez Arias, 2017; Wilson et al., 2009). However, molecular analysis in embryos is limited, because of accessibility to primary material and the challenging temporal resolution. To circumvent these difficulties, over the last few years Embryonic Stem Cells (ESCs) have emerged as a useful model for mammalian development. In the context of axial extension, it has been possible to generate NMPs in vitro from Pluripotent Stem Cells (PSCs) (Edri et al., 2018; Gouti et al., 2014; Gouti et al., 2017; Lippmann et al., 2015; Tsakiridis and Wilson, 2015; Turner et al., 2014). These studies provide large quantities of material and allow the study of details that are difficult to obtain in vivo, particularly the structure and the genetic profile of the NMP population. In these studies, it is important to establish the relationship between the in vitro and the in vivo populations. A recent study aiming to do this and by using an ESCs based protocol, has established some features of an ESC derived NMP population (Gouti et al., 2017).

Here we perform a single cell analysis of different in vitro derived populations comparing them to those in the E8.25 embryo CLE (Ibarra-Soria et al., 2018), where NMPs can be clearly observed (Cambray and Wilson, 2007; Wymeersch et al., 2016). We built a Support Vector Machine (SVM) based on the reference CLE embryo data and used it as a classification model to analyse the different in vitro NMP-like populations and show that while ESCs derived CLE-like populations are heterogeneous and contain few NMP-like cells, EpiSCs derived one, produce a high proportion of cells with the embryo NMP signature. Importantly we find that Epi-CE, the population from which the Epi-NMPs are derived (Edri et al., 2018), contains a node-like population and we show that this population can maintain the expression of *T* in vitro. Our results suggest a sequence of events for the NMPs emergence, which we discuss here.

## Results

**Figure 1.**
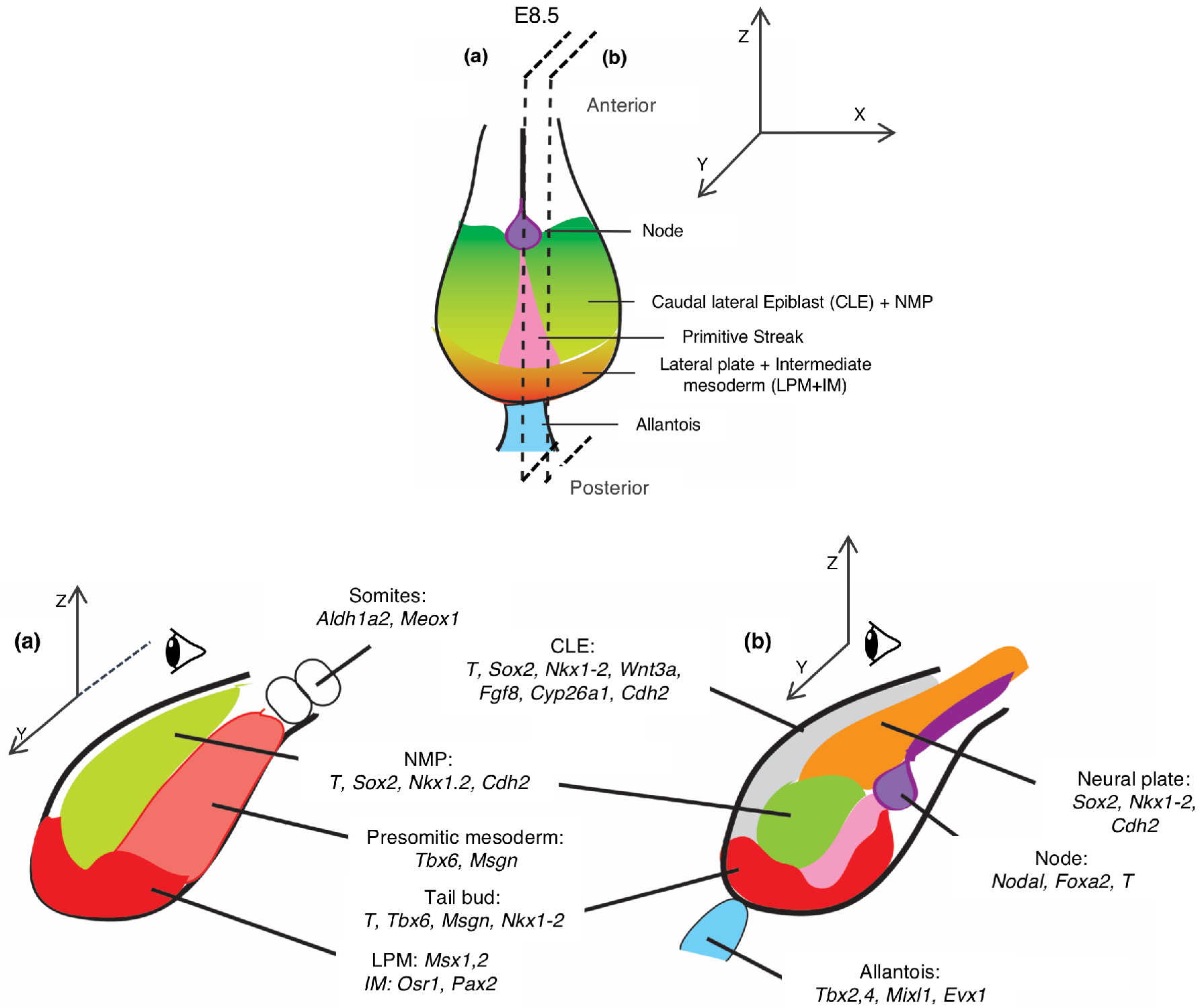
Organization and gene expression patterns in the E8.5 mouse embryo caudal region. Top, ventral view; bottom: lateral (**a**) and medial (**b**) views. The caudal region of the embryo is derived from the posterior epiblast of E7.5 (green in Fig. S1) when the primitive streak (pink) reaches the most distal region of the embryo and the node (purple) appears. This region proliferates and undergoes several morphogenetic events which lead to the organization visible at E8.5 and indicated in the figure. The sources for the outlines shown here can be found in Table S1 and (Edri et al., 2018).

To understand the complexity and identity of the cell populations that emerge when recapitulating NMPs in vitro and how they relate to the embryo CLE, we characterized these populations at a single cell level. We focused our study on the populations that we have described in (Edri et al., 2018) and extracted mRNA from single cells of ES-NMP, (Edri et al., 2018; Turner et al., 2014), Epi-CE and Epi-NMP (Edri et al., 2018), as well as of the *T* expressing cells from the Epi-CE population (Epi-CE-T, Materials and Methods). As a reference for the in vivo population, we used a gene expression data set containing 7,006 cells from E8.25 embryos (Ibarra-Soria et al., 2018). However, rather than using the complete data set, we performed an in silico dissection of the caudal region of the embryo (Fig.1). We selected cells coexpressing *Sox2* and *T* - putative NMPs (Cambray and Wilson, 2007; Henrique et al., 2015; Koch et al., 2017; Tsakiridis et al., 2014; Wymeersch et al., 2016); cells that express *Sox2* and *Nkx1-2* but do not express *T* - preneural progenitors (Henrique et al., 2015; Schubert et al., 1995); and cells that express *T* but not *Sox2*, *Mixl1* or *Bmp4*, which represent mesodermal progenitors and exclude progenitors for the endoderm (*Mixl1*) and the allantois (*Bmp4)* (Dunty et al., 2014; Lawson et al., 1999; Robb et al., 2000; Wolfe and Downs, 2014). We refer to these three population as NMP, preNeuro and preMeso respectively. The extraction process yielded 498 cells that represent the caudal region of the embryo (108 NMP cells, 133 preNeuro cells and 257 preMeso cells).

### In vitro derived populations reflect temporally overlapping embryonic populations

As a first step in our analysis we used the SPRING algorithm (Weinreb et al., 2017) for visualizing high dimensional single cell RNA-seq data (Materials and Methods). This visualization allowed us to obtain a first approximation of the transcriptional complexity of the different samples (Fig. 2 and Supplementary Fig. S2-S6). Each sample occupies a unique position in the dimensionally reduced gene space, with some overlap between the different NMP-like populations. The cells derived from the embryo (blue, Fig. 2) are grouped separately from the in vitro populations and look as an outlier group. Using the reference of the major signature of the CLE gene expression (Fig. 1-2), we observe a spread in the markers expressed by the different populations which can be used to determine their identity.

ES-NMPs appear to be a very heterogeneous population spanning several stages (Fig. 2 and Fig. S2-S6): pluripotency (*Nanog*, *Rex1*, *Sox2*, *Esrrb*, *Fgf4*, Fig. S2), primed epiblast (*Fgf5*, *Otx2* and *Cdh1*, Fig. S2), a later epiblast population that expresses some CLE and NMP markers (Fig. 2 and Fig. S3) as well as cells with a neural identity (Fig. S4) and others with mixed mesodermal characteristic (see Fig. S4). Overlapping with the last population, we notice a group of cells with mixed potential expressing *Mixl1* and *Fgf17* together with *Evx1*, *Hoxb9*, *Oct4* and Wnt genes (Fig. S2 – S6) which might represent the posterior primitive streak population that will give rise to mesendodermal tissue (Dunty et al., 2014; Kojima et al., 2014; Robb et al., 2000; Wolfe and Downs, 2014).The ES-NMPs heterogeneity confirms the conclusion from our previous ensemble study (Edri et al., 2018) that differentiation in the absence of FGF leads to a highly heterogeneous and asynchronous populations with some but few NMPs.

The Epi-NMP population is enriched in cells with expression profiles clearly associated with E8.25/8.5 embryo: expression of *Cyp26a1* and *Cdh2* and absence of *Otx2*, *Oct4*, *Cdh1* and *Fst*, all of which are associated with earlier stages of the embryo (E7.5, Fig. S1 and compare gene expression of E8.25 CLE embryo to Epi-NMP in Fig. 2). In vitro, Epi-NMPs are derived from Epi-CEs (Materials and Methods and (Edri et al., 2018)) which can explain the overlap between the profile of the two populations observed in Figure 2 and how the expression of the different genes indicate a progress in the developmental stage from Epi-CE to Epi-NMP (early epiblast markers in Epi-CE versus CLE markers in Epi-NMP, Fig. 2). We also observe that Epi-NMP, but not Epi-CE, contains a few cells differentiated into mesoderm as highlighted by the expression of *Tbx6*, *Meox1* and *Aldh1a2* (Fig. 1–2). Most surprisingly, we notice that the Epi-CE population, but not Epi-NMP, contains cells coexpressing genes associated with the node e.g. *Nodal*, *Foxa2*, *Ccno*, *Chrd*, *Nog* and *Shh* (Fig. 2 and Fig. S5). A similar population can also be found in the Epi-CE-T and suggests the presence of node like cells in the Epi-CE population. These cells are very reduced in the Epi-NMP population, following the characteristic of the E8.5 CLE (Fig. 1).

**Figure 2.**
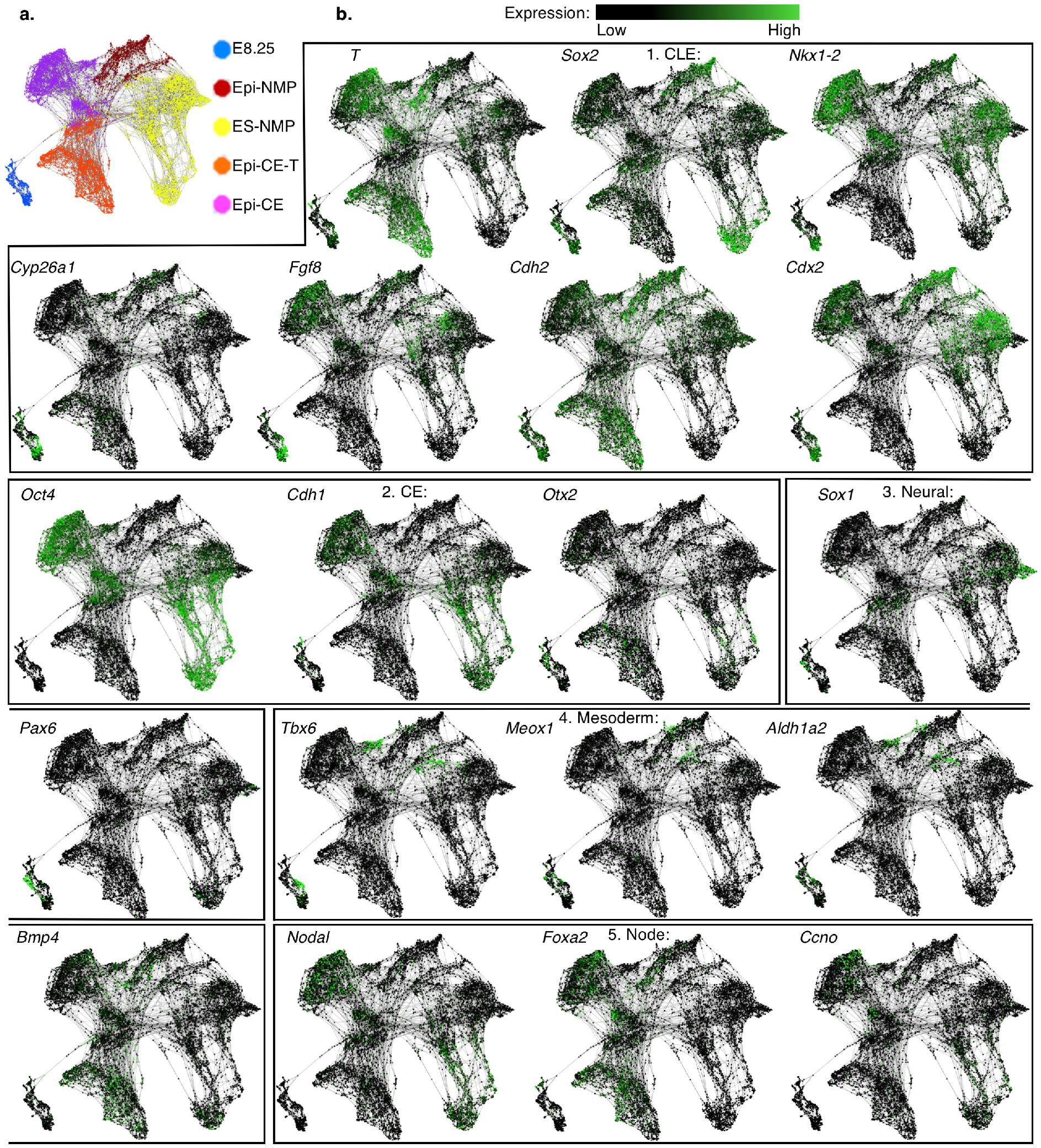
SPRING visualization of gene expression in the in vivo and in vitro populations. **a.** Representation of the different populations. **b.** Expression of chosen marker genes of 1. CLE, 2. CE (E7.5), 3. Neural, 4. Mesoderm and 5. Node along the different samples.

The above observations provide support for our conjecture that that Epi-CE and Epi-NMP correspond to temporally consecutive populations in the embryo, which probably reflect a spectrum between E7.5 (emergence of the node (Davidson and Tam, 2000), Epi-CE) and E8.25/8.5 (Epi-NMP), when NMPs are clearly discernible. The temporal sequence can also be observed in the pattern of Hox genes expression as the Epi-NMP population expresses more posterior Hox genes than the Epi-CE (Fig. S6)

### The NMP landscape in the E8.25 embryo

To interpret the in vitro derived cell populations, we used the caudal cells dissected in silico from the E8.25 embryo to build an SVM pipeline that would enable us to map the NMP-like cells to the in vivo CLE. As a first step, we attempted to identify phenotypically distinct populations amidst the three pools of cells that we had defined based on their pattern of *T*, *Sox2* and *Nkx1-2* expression (Fig. 3a). After processing the single cell data, for both the embryo and the in vitro samples, we found a total of 14,822 genes that can be used for the analysis (Materials and Methods). To provide identifiers genes associated with the CLE region, we based our gene selection on the report from Koch et al. 2017, in which the authors perform an ensemble analysis of the caudal region of the E8.5 embryo based on the levels of *Sox2* and *T*. This work identified 1,402 genes that, together, provide specific signatures for five distinct subpopulations in the caudal end of the embryo: Group 1: axial elongation and trunk development; Group 2: early mesoderm, Group 3: later (committed) mesoderm, Group 4: early neural and Group 5: later (committed) neural ((Koch et al., 2017) and Table S3). Following this study, the genes that are significantly expressed in Group 1, putative NMPs because of the coexpression of *Sox2* and *T*, include in addition genes associated with Group 2 (early mesoderm) and Group 4 (early neural). We used these 1,402 genes and add to them 69 genes which expressed in the decision-making region of the embryo according to the literature (Table S1 and (Edri et al., 2018)), yielding 1,471 genes which were reduced to 1,342 after removal of genes whose mean expression is zero (Table S2). These 1,342 genes were used to cluster the embryo data using a SC3 R package (Kiselev et al., 2017), an algorithm based on k-means clustering (Materials and Methods).

The analysis yielded an optimal number of four clusters in the E8.25 cells (Fig. 3a, Materials and Methods) and 96 marker genes that act as discriminating identifiers of the clusters (Table S3). The top ten marker genes associated with each cluster are visualized in Fig. 3a. Having allocated cells to the 4 clusters based on their gene expression, we looked to see how each of the three functional groups (NMPs candidates, preNeuro and preMeso) that compose the CLE region, occupies each of the clusters.

**Figure 3.**
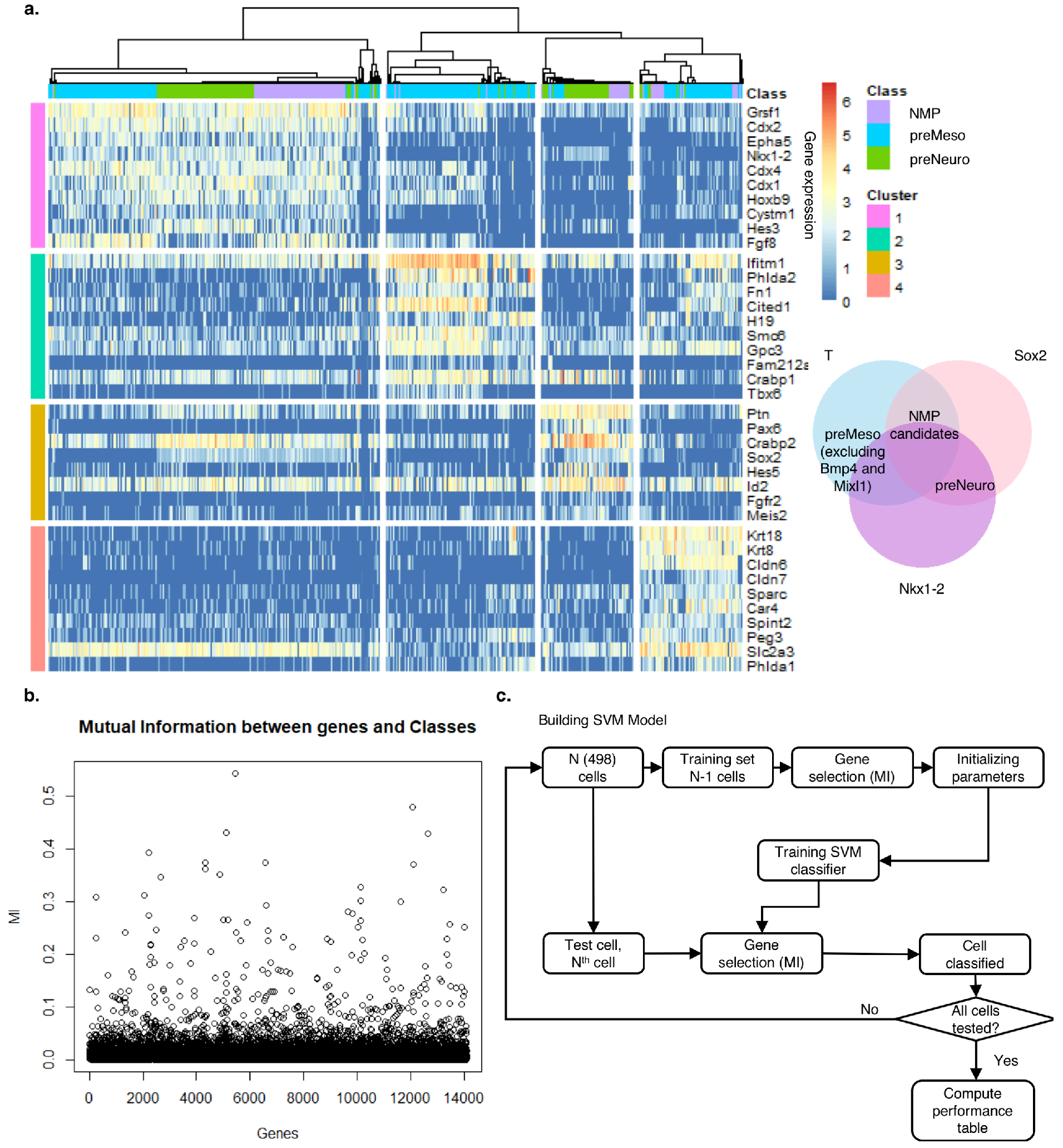
Building SVM based on E8.25 embryo data. **a.** 498 cells representing the CLE and NSB from three E8.25 embryos were dissected in silico and subjected to an unsupervised clustering approach (SC3 R package (Kiselev et al., 2017), Materials and Methods). This yielded 4 clusters and their marker genes: 1) Pink – genes associated with NMPs; 2) Green – mainly mesodermal genes; 3) Dark yellow – genes associated with neural fate, mainly spinal cord; 4) Peach – genes associated with endoderm, mesoderm and extra embryonic tissue (Table S1). **b.** Mutual information (MI) between the genes and the 4 clusters. The informative genes were selected to be above *MI = 0.15*. **c.** Leave one out SVM workflow: an iterative process where each cell is trained and tested (Materials and Methods).

Cluster 1 is a mixed cluster, composed of the three cells categories: NMP candidates, preMeso and preNeuro (Fig. 3a and Table S3); 71% of its 28 marker genes are part of the NMP profile, including *Cdx4*, *Nkx1-2*, *Fgf8* and *Fgf17*(Fig. 3a, Table S3 and (Koch et al., 2017)). Cluster 2 is mainly composed from cells defined as preMeso and the most highly expressed genes in this cluster exhibit a mesodermal affiliation (lateral plate mesoderm (LPM), intermediate mesoderm (IM), PXM and somites, see Table S1) with 91% of the 23 marker genes being mesodermal according to (Koch et al., 2017) (Fig. 3a and Table S3). Cluster 3 is constructed mostly from preNeuro cells and has a neural identity characterized by genes related to the spinal cord and the nervous system. 85% of the 13 marker genes of cluster 3 defined as neural based on the report of (Koch et al., 2017) (Fig. 3a and Table S3). Finally, cluster 4 is mostly composed of preMeso cells and as defined in Koch et al. 2017 34% of the 32 marker genes match to Group 3 (LPM and IM) but with additional genes affiliated to endoderm and IM (Table S1 and Table S3).

Our clustering suggests that cluster 1, which has an NMP signature, might be a more complex population than previously ascertained. It highlights genes like *Nkx1-2*, *Cdx1-4*, *Fgf8*, *Grsf1*, *Epha5* and *Cystm1* associated with NMPs (Cambray and Wilson, 2007; Edri et al., 2018; Gouti et al., 2014; Gouti et al., 2017; Henrique et al., 2015; Koch et al., 2017; Wymeersch et al., 2016), but suggests that rather being a homogeneous population of bipotent cells characterized by the coexpression of *Sox2* and *T*, which only make 29% of this cluster, NMPs might represent a heterogeneous ensemble with additional preMeso (38%) and preNeuro (33%) cells. This analysis raises a question about the differences between the preMeso and preNeuro cells in cluster 1 in comparison to those that are found in clusters 2 and 3. One probable explanation is that cluster 1 encompasses the early committed cells, which are found in the NMP region of the mouse embryo, whereas the others clusters contain more determined cells (Koch et al., 2017). Indeed, cluster 1 includes genes that have previously linked to the NMP profile together with genes that have neural or mesodermal characteristics. Based on the work of Koch et al. 2017 out of the 28 marker genes defining cluster 1, two genes (*Ptk7*and *Fgf8*) are linked to Group 1 (axial elongation and trunk development); 15 genes (*Epha5*, *Nkx1-2*, *Cdx,2,4*, *Cystm1*, *Acot7*, *Stmn2*, *Fgf17*, *Lhpp*, *Mgst1*, *Lix1*, *Hoxc4*, *Ccnjl*, *Sp8* and *Oat*) are linked to Group 4 (early neural) and the rest of the genes are either expressed in the embryo CLE at around E8.5 (*Grsf1*, *Cdx1*, *Hoxb9*, *Hoxc9*, *Wnt5b*), exhibit neural (*Hes3*, *Ncam1*, *Pmaip1*) or mesodermal (*Evx1*, *Hes7*, *Foxb1* which also express in the neural plate) progenitors characteristic (see Table S1 for references).

Having identified a gene based structure for the E8.25 CLE embryo the next step was to build an SVM classifier that will learn the gene profile of the 4 different clusters found in the embryo data. After testing its performance and its stability on the embryo (Fig. 3c, Materials and Methods), the SVM was used to assign cells of the in vitro populations to the 4 classes (clusters) based on their gene expression.

To reduce the number of features (genes) that the SVM needs to learn, we first wanted to identify the informative genes associated with each of the 4 clusters. To do this and to avoid the underrepresentation of genes that were not previously linked to the NMPs, we used the whole set of qualified genes (14,822). The selection of the genes was done by computing the MI between the genes and the 4 clusters (Fig. 3b, Materials and Methods) and resulted in 82 informative genes (Table S4) that were used as features input to the SVM (Fig. 3c, Table S4). 60% of the 82 informative genes are identical to the 96 marker genes of the 4 clusters, whereas 40% of the genes include genes like *T*, *Hoxc8*, *Hoxb8*, *Cdkn1c*, which are expressed in the embryo CLE at E8.25/8.5.

### A comparison between the in vitro and in vivo cell populations

We used the SVM established from the embryo data to explore the structure and nature of the in vitro populations. To do this, we first needed to ensure that the input cells from the in vitro populations, did not contain cells with gene expression patterns on which the SVM had not been trained, as we only want to test the cells with similarity to the E8.25 caudal region (Fig. 3a and step 1 in Fig. 4a). This step resulted in filtering out a higher number of cells from the ES-NMP condition (45%) in comparison to the other conditions (~30%), consistent with the previously noted heterogeneity. Feeding the remaining ‘qualified’ cells to the classifier with only the expression of the 82 informative genes the SVM had been trained on, resulted in the assignment of the probabilities for each cell to be classified to each of the 4 classes (Fig. 4a). In step 6, Figure 4a, only the cells with minimum probability of 0.8 are qualified output cells from the SVM pipeline (Materials and Methods). Again, in this step the highest filter of cells (50%) was observed in the ES-NMP condition compared to the others (~30-35%), suggesting that this condition produce high quantity of cells that do not correspond to the E8.25 embryo CLE. The last step (step 7 Fig. 4a) was to summarize the distribution of the cells of each sample across the 4 classes. Most of the qualified cells (Fig. 4a, Table of step 6) from ES-NMP (84%) and Epi-NMP (73%) were allocated to class 1 (step 7 Fig. 4a), which is associated with the NMPs signature. On the other hand, more than 90% of the qualified cells (Fig. 4a, Table of step 6) from Epi-CE (91%) and Epi-CE-T (97%) were classified to class 4 (step 7 Fig. 4a), which is characterized by the expression of mesodermal and endodermal genes. Class 2 and class 3, which have mesodermal and neural differentiation characteristics, did not attract many cells from the different samples suggesting that the in vitro cells, passed through this pipeline, are not very differentiated.

Figure 4b shows the average expression of the 96 marker genes of the 4 clusters in the in vitro cell populations. This result emphasizes firstly, the fact that the same classes from different samples clustered together, which displays the similarity of the cells from different conditions assigning to the same class. Secondly, that the in vitro cells exhibit the expression of the marker genes of the 4 classes found in the embryo, demonstrating that the SVM pipeline detects the in vitro cells in agreement with the learned embryo cells.

**Figure 4.**
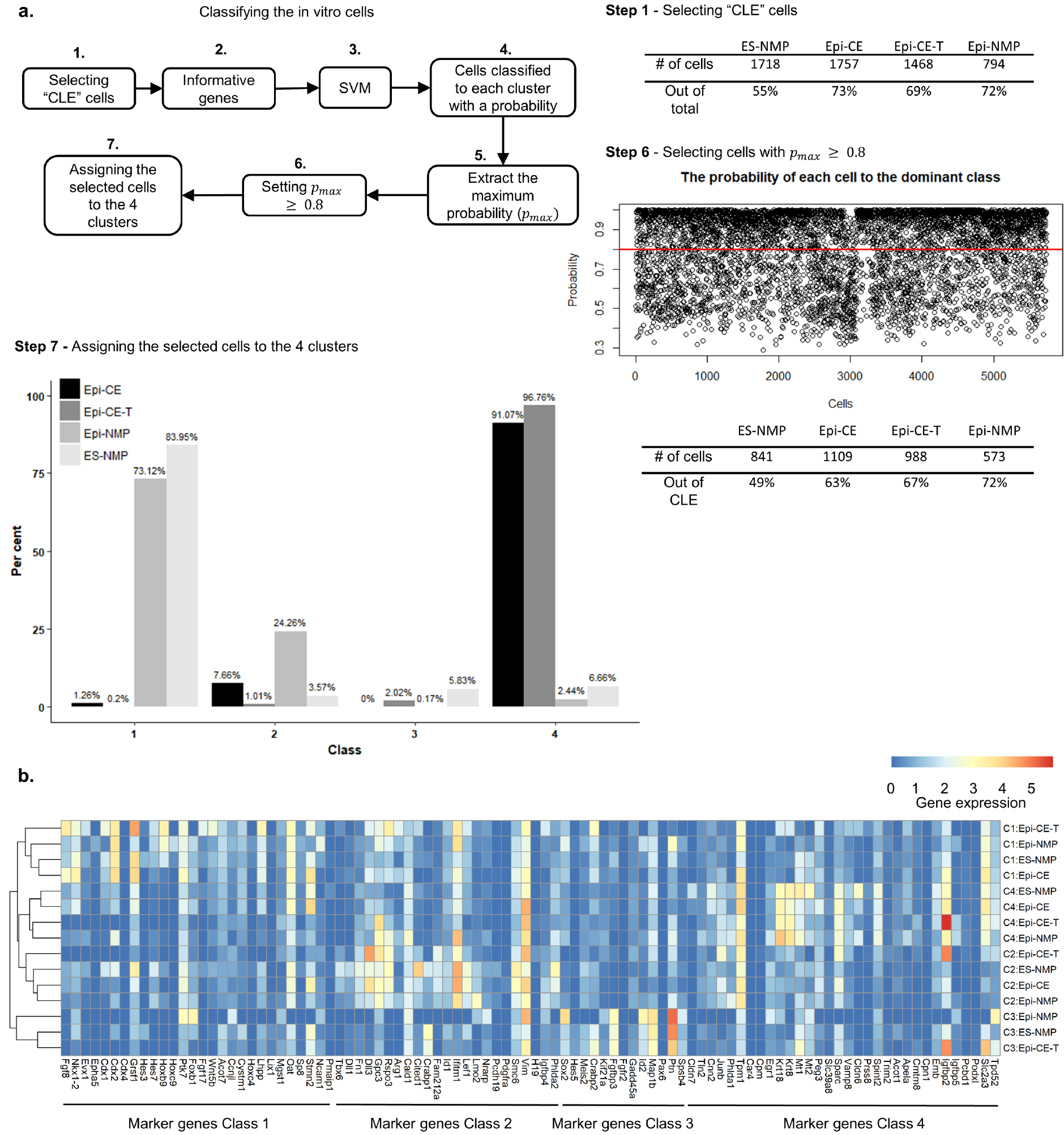
Classifying the in vitro cells using the SVM trained on the embryo data. **a.** Workflow of the classification of the in vitro cells (for details see text and Materials and Methods). **b.** Average expression of the 96 marker genes found in the embryo of each of the 4 classes, in the in vitro samples classified to the 4 classes. Rows of the expression heatmap are hierarchy clustered. Blue-red colour bar indicates the gene expression.

### A node-like population induced in vitro

The finding that Epi-CE and Epi-CE-T, classified mainly to class 4 and that Epi-CE is the origin of Epi-NMP (Materials and Methods, (Edri et al., 2018)) led us to investigate further the identity of cluster 4. As a first step, we arranged all the qualified cells out of the SVM pipeline (Fig. 4a, Table of step 6) into a pseudotime ordering using TSCAN, a Biocounductor R package version 1.16.0, (Ji and Ji, 2017) (Materials and Methods). This analysis revealed that class 4 cells (red cells in Fig. 5a) are split into 2 pseudotime ranges, with class 1 cells (blue cells in Fig. 5a) forming a bridge between these two classes. This result lends support to the fact that Epi-NMP cells (mainly classified to class 1) are derived from Epi-CE (class 4 mainly composed from Epi-CE and Epi-CE-T). It also raises the existence of two different populations in Epi-CE. When exploring the highly expressed genes that define the 2 pseudotime ranges of class 4 (Fig. 5a, Materials and Methods and Table S5), we observed that the later range is defined by genes associated with rapidly dividing cells, whereas the early one doesn’t show this enrichment (Fig. 5a). This observation suggests the existence, in class 4, of a group of cells in a phase of large expansion.

The presence of endodermal and mesodermal markers in class 4 is surprising as it suggests the existence of a cell type in the embryo caudal region, that would be associated with these germ layers. One structure that could fit this criterion is the node (Blum et al., 2007; Lee and Anderson, 2008; Martinez Arias and Steventon, 2018), a structure that appears at E7.5, contains the progenitors of the axial mesoderm (Beddington, 1982; McGrew et al., 2008; Tam and Beddington, 1987) and has been associated with the NMPs (Albors and Storey, 2016; Garriock et al., 2015; Henrique et al., 2015; Wymeersch et al., 2016). Thus, we considered the possibility that class 4 contains node cells.

At a very coarse level, the node can be identified as cells expressing combinations of three genes; *Foxa2*, *Nodal* and *T* (Fig. 5b (Davidson and Tam, 2000; Jeong and Epstein, 2003a; Lee and Anderson, 2008; Shiratori and Hamada, 2006)). Applying this coarse definition, we detected node-like cells in our in vitro samples with a very high representation in class 4 (Fig. 5c). The allocation of a node identity to cells in class 4 is not a bias of the sample size, as a statistical test controlling the size of the classes yielded that class 4 has the highest proportion of node-like cells is statistically significant (*p* - *value* < 0.001, Materials and Methods). To further test this coarse identification of node-like cells, we gathered a list of additional genes associated with the structure and function of the node e.g., *Shh*, *Ccno* and *Chrd* (Davidson and Tam, 2000; Funk et al., 2015; Jeong and Epstein, 2003a; Lee and Anderson, 2008; Shiratori and Hamada, 2006; Tam and Behringer, 1997) and tested for their expression in class 4 as can be seen in Figure 5d.

Having identified node-like cells in our in vitro populations we thought we could use the dynamic changes in this region of the embryo to stage our in vitro populations. For example, at the time of its appearance the node expresses *Oct4* and *Otx2* however by E8.0-8.5 the expression of these genes have disappeared from the node (Cajal et al., 2012; Downs, 2008). The expression of *Oct4* is particularly diagnostic for this transition.

**Figure 5.**
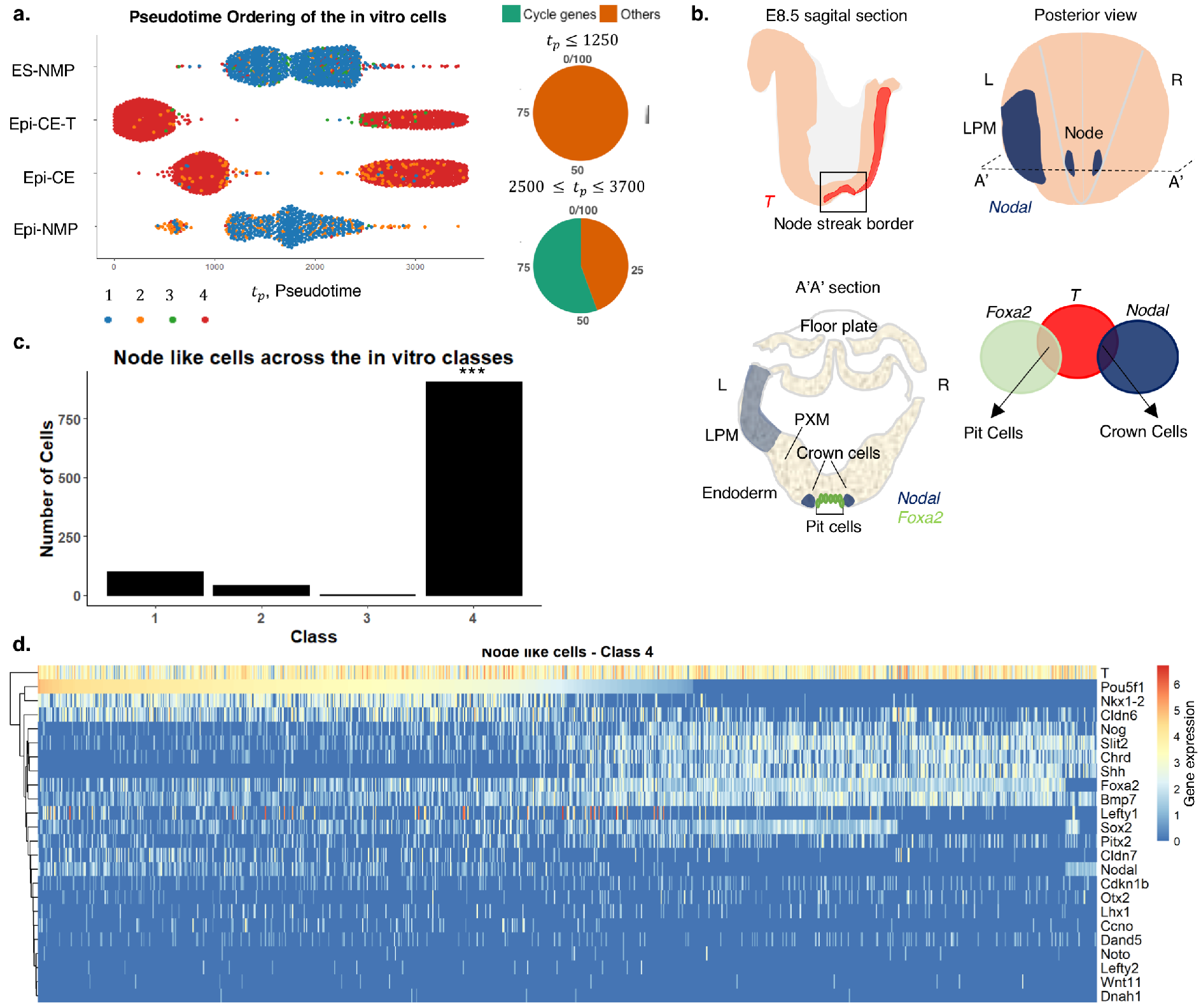
Class 4 contains node-like cells. **a.** Pseudotemporal order of the in vitro cells that were classified to the 4 classes. Class 4 is divided to 2 pseudotime ranges: the highly expressed genes in the later range contains 55% of cycling genes whereas the early one doesn’t contain any cycling genes (Materials and Methods), **b**. E8.5 mouse embryo node: illustration of a sagittal view of the embryo shows the expression of *T*(red) in the NSB (Tsakiridis et al., 2014). Posterior view of the embryo exhibits the expression of *Nodal* (blue) in the node and in the LPM (Shiratori and Hamada, 2006) and its left (L) right (R) asymmetry. A transverse section (A’A’) reveals the pit and crown cells of the node, PXM, LPM, endoderm and the prospective floor plate. The expression of *Nodal* and *Foxa2* is indicated in blue and green respectively. The pit cells coexpress *T* and *Foxa2* and the crown cells express *Nodal* and *T* (Davidson and Tam, 2000; Jeong and Epstein, 2003a; Lee and Anderson, 2008; Shiratori and Hamada, 2006). **c.** The distribution of the node-like cells amongst the 4 classes: significant higher number of the node-like cells are found in class 4 in comparison to the other classes. **d.** Gene expression heatmap of chosen node genes in class 4. The genes are hierarchically clustered and the cells are ordered in accordance with the decreasing expression of *Oct4* (*Pou5f1*). Gene expression, which is defined as *log*_2_(*CPM* + 1) (Materials and Methods), is indicated by the blue-red colour bar.

Ordering node-like cells of class 4 (Fig. 5d) from high to low *Oct4* expression, reveals additional patterns of gene expression that confirm the presence of a node-like population in the in vitro class 4 associated with *Oct4* expression. Cells with decreasing levels of *Oct4* display increasing levels of genes associated with the node: *Foxa2*, *Bmp7*, *Noggin*, *Chrd*, *Slit* and significantly *Shh* (Fig. 5d and (Davidson and Tam, 2000)). Within the cells expressing low or no *Oct4*, we observe a further division based on *Sox2* expression: while all cells express node genes, some of them express *Sox2* and some don’t. The ventral most region of the neural plate is called the floor plate (Fig. 5b) and shares many of the pattern of gene expression of the node (Jeong and Epstein, 2003b; Wood and Episkopou, 1999). These results suggest that our experiment not only yields node-like cells (*Sox2* negative) but also floor plate precursors (*Sox2* positive).

**Table 1.** The distribution of the node-like cells in the 2 pseudotime ranges of class 4 of the in vitro cells.

|  | Pseudotime range | Class 4 | Node-like cells | Out of class 4 |
| --- | --- | --- | --- | --- |
| Time group 1 | $tp \leq 1250$ | 1050 | 433 | 41% |
| Time group 2 | $2500 \leq tp \leq 3700$ | 916 | 442 | 48% |

Moreover, we checked the proportion of the node-like cells in the 2 pseusotime ranges of class 4, as shown in Table 1, however no difference was found. This result, supports our hypothesis that these two populations are very similar, however one of them represents an amplifying population versus the second one which is more stable in terms of size.

In the in silco E8.25 embryo CLE we found 38 node-like cells (Fig. 6a-b), 30 of which were mapped to the embryo class 4 (Fig. 6a). When comparing the node-like cells in class 4 of the embryo to those of the in vitro cells, some notable differences become apparent (Fig. 6b). For example *Oct4*, which is off in the embryo cells. Since Epi-NMPs express very few node genes (Fig. 4a step 7 and Fig. 6c) and no *Oct4*, first supports our previous assertion that it has the closest relationship to the E8.25 CLE region, second that the node-like cells are lost in the transition between Epi-CE and Epi-NMP (Fig. 6c) and that the Epi-CE cells represent a developmentally earlier cell state than the Epi-NMPs.

### An in vitro functional test of the in vitro induced node-like population

**Figure 6.**
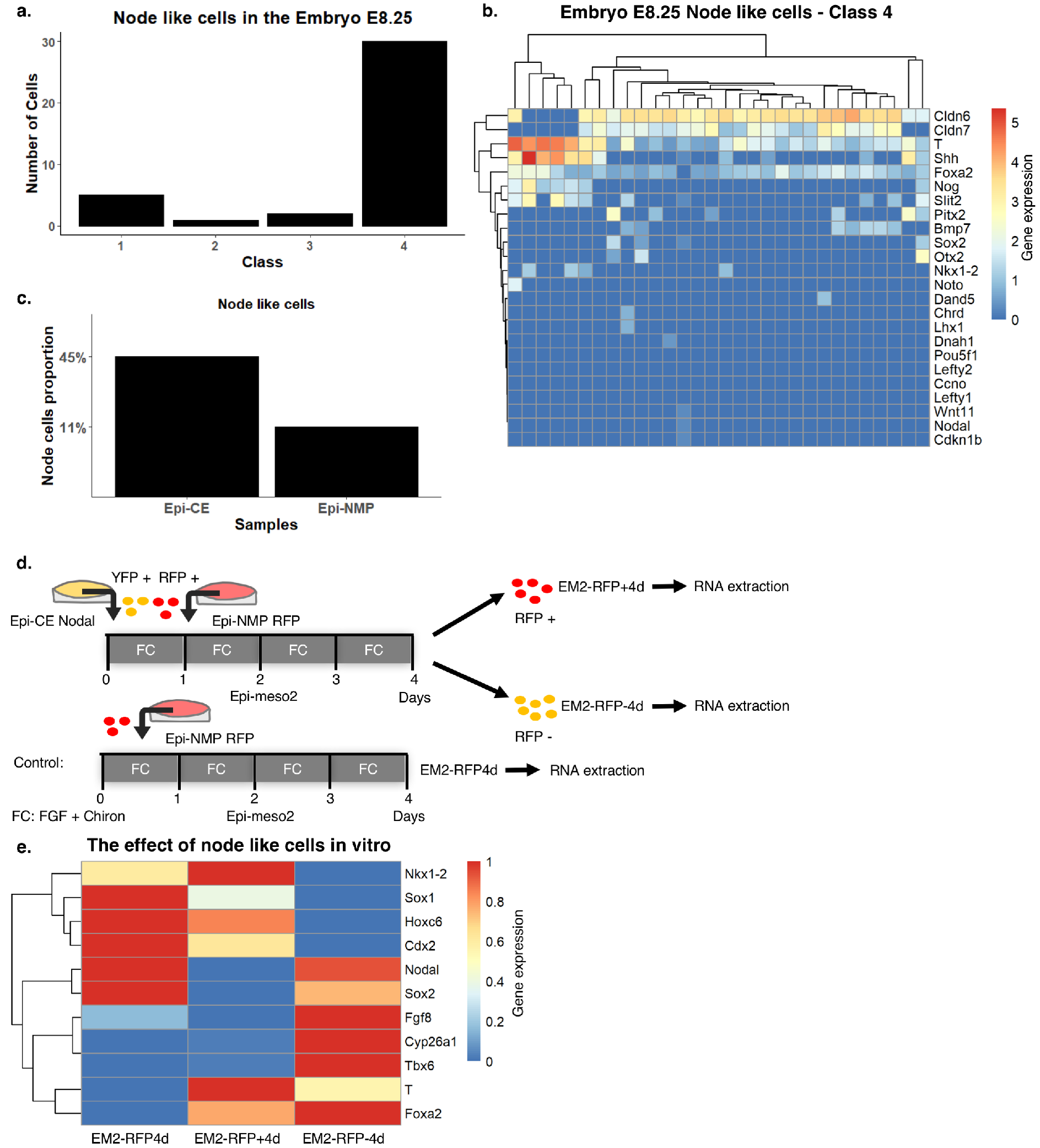
Node cells are needed to maintain the NMPs. **a.** The distribution of the node cells amongst the 4 classes in the embryo. **b.** Expression of chosen node genes in the embryo class 4. Genes and cells are hierarchically clustered. Gene expression which is defined as *log*_2_(*CPM* + 1), is indicated by the blue-red colour bar. **c.** Proportion of node-like cells in the Epi-CE and Epi-NMP samples. **d.** YFP positive cells of Epi-CE Nodal sample composed of Nodal::YFP cells and RFP positive cells of Epi-NMP RFP sample composed of Ubiquitin::Tomato cells, were used to make Epi-meso2 mixture (Materials and Methods). This mixture was grown for 4 days then the cells were sorted based on their RFP fluorescence: RFP positive cells (sample named EM2-RFP+4d) and RFP negative cells (sample named EM2-RFP-4d). the control sample is Epi-NMP RFP sample composed of 100% Ubiquitin::Tomato cells that was cultured for 4 days in FGF and Chiron to make Epi-meso2 (sample named EM2-RFP4d). The sorted cells and the control sample were quantified for their mRNA of a chosen set of genes using RT-qPCR technique. e. Expression heatmap of 11 genes, obtained by RT-qPCR, in cells grown in the 3 conditions, as indicated in Fig. 6d. The normalized expression of each gene to the housekeeping gene *Ppia* was scaled between 0 and 1 across the different conditions. Gene expression is indicated by the blue-red colour bar.

Previously we showed that the Epi-NMP population has a limited but clear self renewing ability in culture when exposed to FGF and Chiron (Edri et al., 2018). These cells maintain *T* and *Sox2* expression for at least two passages (Epi-NMP, Epi-meso2, Epi-meso3…) though, over time, the levels of NMP markers go down and the cells exhibit a slow increase in the expression of differentiation genes associated with neural fates (Edri et al., 2018). In the embryo, the self renewing population also decreases with time and this is associated with the node disappearance (Steventon and Martinez Arias, 2017; Wymeersch et al., 2016). Thus, we considered that, in our in vitro system, the loss of *T* might be associated with the loss of node-like cells. To test this, we added node-like cells from Epi-CE to Epi-NMP and passaged the mixed sample to make Epi-meso2, then we checked whether the addition of node-like cells could maintain the levels of *T* expression.

The experiment is described in Figure 6d: as a source of NMP-like cells we used a Ubiquitin::tomato cell line and as a source of node cells, a Nodal::YFP cell line. Both were cultured to produce Epi-CE: Epi-CE RFP (from Ubiquitin::tomato cell line) and Epi-CE Nodal (from Nodal::YFP cell line). The Epi-CE RFP were further grown to make Epi-NMP (Epi-NMP RFP). After two days of culturing Epi-NMP RFP, we plated a mixture that equally consists of Epi-NMP RFP positive cells and Epi-CE Nodal positive YFP cells (Fig. 6d, Fig. S8a-b, Materials and Methods). The mixture (Epi-meso2) was cultured for 4 days (Fig. 6d) until sorting the cells to RFP positive (sample named: EM2-RFP+4d, contains only the Ubiquitin::tomato cells) and RFP negative (sample named: EM2-RFP-4d, contains Nodal::YFP cells and might contain Ubiquitin::tomato cells that didn’t express RFP, Fig. S8c and Materials and Methods). These populations of cells were compared to the EM2-RFP4d, which are only Epi-NMP RFP cells cultured for 4 days to make Epi-meso2 (Fig. 6d, Materials and Methods).

Addition of node-like cells to the Epi-NMP population elevates the level of *T* and *Foxa2*, maintains the expression of *Cdx2* and *Nkx1-2* and decreases the level of neural fate markers: *Sox2* and *Sox1* (Figure 6e and Figure S9 EM2-RFP4d versus EM2-RFP+4d). In addition, there is no much difference in the expression of *Tbx6*, *Hoxc6*, *Fgf8* and *Cyp26a1*, when comparing those genes between EM2-RFP4d and EM2-RFP+4d.

This result, aligned with what we previously showed (Edri et al., 2018), suggests that node like cells are necessary to maintain the relative levels of *Sox2* and *T* and buffer the tendency that the Epi-NMPs have towards the neural fate when passaging them in culture.

## Discussion

Over the last few years, ESCs have emerged as a useful experimental system to study mammalian development. While they are no substitute for the embryo, they have some advantages when addressing processes that happen early in development, when material and experimental accessibility are scarce. However, their validation as an experimental system depends on showing how they relate to events in the embryo. Here we have used mouse PSCs to analyse the origin and structure of NMPs, a bipotent population that is thought to give rise to the spinal cord and the paraxial mesoderm. As an important reference for our study we have used a single cell data set from E8.25 embryos, the stage at which NMPs are first distinguishable.

Analysis of NMPs derived from PSCs suggests that different protocols produce heterogeneous populations in terms of gene expression. To gain insights into these heterogeneities and their complex origins, we have performed a single cell transcriptomics analysis of the different populations. As a reference, we have used data from E8.25 embryos out of which we have dissected in silico the CLE/NSB region based on *T*, *Sox2* and *Nkx1-2* expression patterns, as cells that express these genes are often identified as NMPs. Our results suggest that, by this stage, these cells are distinct from those in the pluripotent epiblast. The transition between the states appears to be associated with the expression loss of *Cdh1*, *Oct4*, *Fst* and *Otx2* and the gain of *Cdh2* and *Cyp26a1* amongst others (Fig.1 and Fig.S1). Our results contrast with those of a recent study which allocated expression of *Cdh1* and *Oct4* to NMPs at E8.5. Analysis of published gene expression patterns (Fig.1, Fig.S1 and Table S1) supports our conclusions that these markers are associated with the pluripotent epiblast. It might be that changes in the transcription of these genes happens abruptly at around E8.25 and that there is a difficulty in staging the embryos. The transition from pluripotent epiblast to the bipotent cells in the CLE/NSB region can be detected in our vitro samples as represented by the transition from Epi-CE to Epi-NMP (Fig. 2 and Fig. S2-S6).

Using a clustering algorithm, we identified four populations in the embryo: class 1 with NMP signature; class 2 with mesodermal signature; class 3 with neural signature and class 4 with extraembryonic, endoderm and IM signature. Class 1 contains cells coexpressing *Sox2* and *T* and cells in a pre neural or mesodermal state, i.e. not all of them coexpress *Sox2* and *T*. This emphasizes the notion that NMPs should coexpress *Sox2* and *T* is not a valid definition, or at least it is not an absolute condition for defining NMPs. This also raises the possibility that an NMP population is not only a collection of poised *Sox2* and *T* coexpressing cells (Gouti et al., 2014; Gouti et al., 2017; Turner et al., 2014), but rather a heterogeneous population of poised and early differentiated cells, perhaps in dynamic equilibrium.

The structure of the E8.25 caudal region was used as a reference for the analysis of the in vitro derived populations. To do this, we used the four classes derived from the embryo data to build an SVM classification model that allowed us to allocate cells from the different protocols to our reference. We find that the ESC based protocol contains few cells allocated to the E8.25 embryo CLE, but that the EpiSCs samples are enriched. Furthermore, we find that Epi-NMP cells, which are derived from Epi-CE (Materials and Methods) contain the most E8.25 CLE-like cells (>70% of the selected cells, Fig. 4a) and most of them map to class 1. Furthermore, we find many E8.25 CLE-like cells in the Epi-CE population (>60% of the selected cells, Fig. 4a), but in contrast with Epi-NMPs, these cells predominantly map to class 4. Interestingly, very few cells of the Epi-CE descendant, Epi-NMP, map to class 4. A detailed analysis of class 4 reveals that it has a large representation of node-like cells and, interestingly, of the floor plate. The floor plate in the embryo shares many features with the node and its main derivative, the notochord. This allocation is confirmed by the identification of node-like cells in the embryo reference data.

The representation of cells from two different sequentially induced in vitro populations to one embryonic stage is, at first sight surprising, however we believe that there is an explanation. The CLE at E8.25 is derived from an earlier caudal region, at E7.5, whose most prominent feature is the node, that is maintained until E9.0. Thus, at E8.25 the embryo does have a signature of an early stage in the node. The representation of node cells in Epi-CE but not much in its progeny, Epi-NMP, suggests that, in adherent culture the conditions are not conducive to the maintenance of the node. What we find interesting, given the relationship between Epi-CE and Epi-NMP, is the presence of NMP-like cells in the Epi-NMP population. This would suggest that in the embryo there is a very close relationship between the node and the NMPs emergence, something that has been suggested before (Albors and Storey, 2016; Garriock et al., 2015; Henrique et al., 2015; Wymeersch et al., 2016), and our in vitro system recapitulates this relationship.

We find two interesting features of the relationship between these two populations. The first one is the observation that in the Epi-CE population there is a subpopulation in a high proliferative state and the second one is the relationship we have observed between the node and the maintenance of the *T* and *Sox2* expression ratio. These observations led us to suggest that, in the embryo, NMP population arise early in development, near the node and that the node plays a role in its maintenance and amplification at that early stage. A need for amplification of the initial NMPs pool could be likely explained by the size of the primordia relatively to the size of the tissue that needs to be generated. It is not clear how the node mediates this function but an interaction between BMP and Nodal (Edri et al., 2018) might be important. A relationship between the node and axial elongation can be gauged from the effect of mutations in which the node is absent. This leads to a loss of *T* expression in the caudal region of the embryo and severe truncations (Ang and Rossant, 1994; Davidson and Tam, 2000; Weinstein et al., 1994). The effect of *Oct4* might be important as we observe a clear transition in the behaviour of the in vitro populations whether they express *Oct4* (Epi-CE) or not (Epi-NMP). This might correspond to the proliferative amplification phase and the start of the differentiation phase of the NMPs. In this regard, it might be that *Oct4* creates a molecular context for *Sox2*; as long both are expressed the cells in the epiblast are multipotent and only when *Oct4* is downregulated, *Sox2* becomes engaged in neural differentiation. It will be interesting to test this hypothesis.

## Acknowledgements

We would like to thank Bertie Gottgens for sharing the embryo data before publication, Meritxell Vinyoles for helping in experimental design and insightful discussion and to James Briscoe, Valerie Wilson and Ben Steventon for insightful discussions.

## Materials and Methods

### ESC Culture and routine cell culture

E14-Tg2A, Bra::GFP (Fehling et al., 2003), Nodal::YFP (Papanayotou et al., 2014) and Sox17::GFP Ubiquitin::Tomato (Niakan et al., 2010) mouse ESCs were cultured in tissue-culture plastic flasks coated with 0.1% gelatine in PBS (with Calcium and Magnesium), filled with GMEM (Gibco, UK) supplemented with non-essential amino acids, sodium pyruvate, GlutaMAX™, β-mercaptoethanol, foetal bovine serum and LIF. Cell medium was changed daily and cells passaged every other day.

The differentiation protocols are as the following:

#### ES-NMP

E14-Tg2A cells were plated at a density of 4.44×10^3^ cells/cm^2^ in a 0.1% gelatine coated flask with a base medium of N2B27 (NDiff 227, Takara Bio) for 2 days. After 48hr N2B27 is supplemented with 3μM of CHIR99021 (Chiron 10mM, Tocris Biosciences) for additional 24hr, which are in total 72hr.

#### EpiSCs

E14-Tg2A or Bra:GFP were grown on a 0.5% Plasma Fibronectin (FCOLO, 1mg/ml, Temecula) in PBS (with Calcium and Magnesium) coated culture flask with N2B27 supplemented with 12ng/ml FGF2 (R&D systems, 50*μ*g/ml) and 25ng/ml Activin A (Stem Cells Institute 100μg/ml), known as Epi-media, for at least 4 passages. These cells considered as EpiSCs. Those cells can be tested to be EpiSC by seeding them in a colony assay density (67 cells/cm^2^) in restricted medium (2i: N2B27 supplemented with 3μM Chiron and 1 μM PD0325901 (PD03, Tocris Biosciences, 10mM)), resulting in no growth of cells, ensuring that the cells are no longer in the naïve pluripotent state and they moved on to the prime pluripotent state (data are not shown).

#### Epi-CE and Epi-CE-T

EpiSCs were plated at a 5×10^4^ cells/cm^2^ density in a 0.5% Fibronectin pre-coated flask with Epi-media for the first day. Day 2 is followed by increasing the concentration of FGF2 to 20ng/ml in the base medium of N2B27 and removing Activin A. On day 3, N2B27 is supplemented with 3μM Chiron which is added to the 20ng/ml FGF2. After 72hr those cells known as the Epi-CE. This protocol is a variation of one that has been used to derive NMP-like cells from human ESCs (Lippmann et al., 2015). Epi-CE-T were cultured from Bra:GFP cell line at the same way as Epi-CE with the modification that after 72 hours the cells were sorted for positive GFP cells only.

#### Epi-NMP

Epi-CE cells were detached from the culture flask using Accutase (BioLegend 0.5Mm) and seeded on a flask coated with 0.5% Fibronectin at a dense of 5×10^4^ cells/cm^2^. The cells were grown for 2 days in N2B27 supplemented with 20ng/ml FGF2 and 3μM Chiron.

### Single cells transcriptomic

10× Genomics single cell transcriptomic service was used to sequence our 4 different samples. 8,700 cells from each sample were loaded into the 10× Chromium system. The preparation of the libraries and the Illumina sequencing (HiSeq 4000) was done by the Cambridge 10X genomics services. Cell Ranger version 1.3.1 (10× Genomics) was used to process raw sequencing data and the Seurat R package version 2.0 (Butler and Satija, 2017; Macosko et al., 2015) was used to read the data from Cell Ranger to R and build the expression matrix. Gene expression was quantified by UMI counts. The final output was a matrix of genes versus cells, utilized for further analysis.

### Embryo data

In this work, we used the published transcriptomic single cell data from 3 mouse embryos females and males at E8.25 (Ibarra-Soria et al., 2018) including their extra-embryonic tissues. These embryos were dissociated to single cells and processed on a 10X microfluidic chip. The resulting libraries were sequenced on an Illumina HiSeq 2500, subsequent in total 7,006 cells out of which 4,706 are male cells and 2,300 are female cells.

### Single cell data clean up and quality control

Using Scater package in R (McCarthy et al., 2017), the expression matrix was cleaned in the 4 following aspects: 1) UMI counts – drawing the histogram of the RNA UMI total counts per cell, allowed us to set a threshold of above 8,000 UMI counts in a cell, ensuring a sufficient sequencing depth for each cell; 2) detected genes – from the histogram of total detected genes in a cell we set a threshold of above 2,500 unique genes in a cell, ensuring the reads are distributed across the transcriptome; 3) mitochondrial genes expression - plotting the per cent of mitochondrial genes counts in a cell versus the total detected genes in a cell, allowed us to set a threshold of 20%, ensuring the cells to be further analyse are not likely to be dead or stressed; 4) Gene filtering – undetectable genes were filtered out by setting a threshold of having at least two cells containing more than 1 UMI of a gene. After the clean up the number of cells and total genes are presented in Table 2.

**Table 2.** The number of cells in each sample and the total number of detected genes after single cell data clean up.

| **Sample** | **Total cells** | **Total genes** |
| --- | --- | --- |
| ES-NMP | 3,133 | 14,822 |
| Epi-CE | 2,404 | 14,822 |
| Epi-CE-T | 2,135 | 14,822 |
| Epi-NMP | 1,108 | 14,822 |
| Embryo E8.25 | 4,183 | 14,822 |

The UMI count normalization, which is necessary to make accurate comparison of gene expression between samples, was done by first scaling the counts of each gene in a cell to the total counts in that cell per million counts (known as counts per million, CPM). Then the log_2_(CPM+1) was calculated for each gene, this is the normalized gene expression (the 1 was added to the CPM to keep zero counts as zero in the binary logarithm scale).

**Visualization of the single cell data using SPRING** (Weinreb et al., 2017) SPRING is composed of the following steps:

1. Filters cells with fewer 1000 UMIs.
2. Filters genes with average expression > 0.05 UMIs/cell.
3. Normalize gene expression that every cell has the same total reads.
4. filters for genes with average expression > 0.1 and Fano factor > 3.
5. Z-score normalize expression and reduces dimensionality to a 20 dim PCA space.
6. Compute distance matrix and output k-nearest-neighbour (knn) graph.

This visualization exhibit how cells are positioned in high-dimension with respect to one another.

### Clustering the embryo cells

The dissection of CLE in silco from the whole mouse embryo was done by selecting cells that coexpress *Sox2* and T; cells that express *Sox2* and *Nkx1-2* and don’t express *T* and cells that express *T* and don’t express *Sox2*, *Mixl1* and *Bmp4* (see text). Before clustering the embryo CLE cells, a selection of genes was carried out to guide this action. The selection was made to get the focus on the caudal region of the embryo and, importantly, to avoid biases towards clustering results led by genes associated with different processes or regions; for example, the embryo data is a mixture of male and female embryos and in this situation, *Xist* expression leads to clusters of female and males (unpublished observation). The genes that were selected for our analysis were a total of 1,402 genes reported by Koch et al. 2017 in a study of the NMPs and the caudal region of the embryo (Koch et al., 2017). To this list further genes were added due to their association with the CLE region of the E8.5 embryo (Edri et al., 2018), reaching a total of 1,471 genes. From this list, genes with zero mean expression were removed, yielding a total of 1,342 genes for analysis (Table S2). Clustering was performed with the Cell Consensus Clustering (SC3) package in R (Kiselev et al., 2017) with the following steps:.

1. Gene filter – filtering genes that are either expressed in less than *6%* of the cells (rare genes) or expressed in at least 94% of cells (ubiquitous genes).
2. Distance matrices calculations – distances between the cells are calculated using the Euclidean, Pearson and Spearman matrices.
3. Transformations – All distance matrices are then transformed using either principle component analysis or by calculating the eigenvectors of the associated graph Laplacian.
4. k-means – k-means clustering is performed on the first set of eigenvectors of the transformed distance matrices. The number of clusters k is set by the user.
5. Consensus clustering – a binary similarity matrix is constructed for each individual clustering result from the corresponding cell labels obtained in the previous step: if two cells belong to the same cluster, their similarity is 1; otherwise the similarity is 0. A consensus matrix is calculated by averaging all similarity matrices of the individual clustering results. The resulting consensus matrix is clustered using hierarchical clustering.

The clustering of the embryo cells was done between k = 2 and k = 8. The Consensus matrices for the different k are shown in Figure S7. The averaged Silhouette width for each clustering results is summarized in the table below:

**Table 3.** The averaged Silhouette width of the consensus matrices obtained for clusters ranging from k = 2 to k = 8.

|  | k = 2 | k = 3 | k = 4 | k = 5 | k = 6 | k = 7 | k = 8 |
| --- | --- | --- | --- | --- | --- | --- | --- |
| **Averaged Silhouette width** | 0.8 | 0.9 | 0.85 | 0.78 | 0.77 | 0.77 | 0.72 |

The silhouette is a quantitative measure represents the consensus matrix diagonally. An average silhouette width, which is calculated as the weighted average between the silhouette values of each cluster, varies from 0 to 1 and the closest it is to 1 the better the clustering is for that value of k. From the consensus matrices on Figure S7 and from the averaged silhouette width in Table 3, we estimated that the optimal number of clusters could be k = 3 or k = 4. For k = 3, the 3 clusters are: a mixed cluster – containing cells from all the 3 categories: NMP candidates, preNeuro and preMeso; and the two other, mainly composed from preMeso cells (Fig. S7). For k = 4 the clusters are: a mixed cluster composed from all the three cells’ categories, 2 others which are mainly composed of cells with a mesodermal identity and a forth one which is mainly constructed from neural oriented cells (Fig. S7). We decided to continue to downstream analysis with k = 4 because of the appearance of a clear neural along with mesodermal clusters. K = 4 ensures a representation for all the three cells’ categories: NMP candidates, mesodermal and neural cells.

### Marker genes

Using the SC3 package in R (Kiselev et al., 2017) 96 marker genes were identified for the 4 obtained clusters (see Table S3). Marker genes are defined as genes that are highly expressed in only one of the clusters and can lead to the segregation of one cluster from the rest. Finding the marker genes are according to the following steps as was explained in (Kiselev et al., 2017):

1. Constructing a binary classifier for each gene based on comparing the mean expression values across the clusters.
2. Calculating the classifier prediction by comparing the gene expression ranks across clusters.
3. Quantify the accuracy of the prediction by calculating for each gene the area under the receiver operating characteristic (ROC) curve (true positive rate versus false positive rate).
4. Calculating the p-value for each gene by using the Wilcoxon signed rank test: comparing the gene ranks in the cluster with the highest mean expression with all others.
5. Setting a threshold for the area under the ROC curve and the p-value to determine the marker genes.

The genes with the area under the ROC curve > 0.65 and with the p-value < 0.01 are defined as marker genes. The top 10 marker genes of each cluster are visualized in Fig. 3a.

### Mutual information between genes and classes

After identifying the 4 different clusters in the in silico CLE embryo data, the downstream analysis was constructed with the whole set of qualified genes (14,822) rather than with the genes restricted to CLE (1,342). This step was performed to avoid an underrepresentation of genes that were not previously linked to the NMPs. However, there is a need for dimensionality reduction to elucidate the data and to feasibly reduce computer calculation time. Here, similar to the work of Vanitha et al., 2015 (Vanitha et al., 2015), we used a mutual information (MI) technique (Battiti, 1994) to select the informative genes related to the 4 clusters. The steps of computing the MI between the clusters (denoted as Y) and genes (denoted as X) are the following:

1. Calculating the clusters entropy:

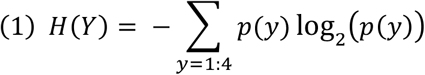

Where *p*(*y*) is the probability of each cluster *y* = 1,2,3,4, which is computed based on the distribution of the 4 clusters in the embryo data.
2. Discretization the gene expression values into ten bins and calculating the conditional entropy *H*(*Y*|*X*) as the following:

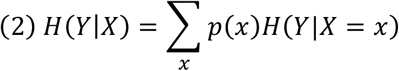

Where *p*(*x*) is the probability of the discretize expression values of a gene across the cells population and *H*(*Y*|= *x*) is the clusters entropy given a specific gene expression value, calculated following Equation 1.
3. Computing the MI between the clusters and each gene is accordant the below equation:

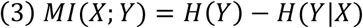
4. Setting a threshold of the MI of all the genes and selecting the informative genes above this value (Fig. 3b and manuscript in preparation) to train the SVM.

**Table 4.** Performance of the SVM with setting different threshold of the MI value

|  | Number of<br>informative<br>genes | Correctly<br>classified | Misclassified | Error rate |
| --- | --- | --- | --- | --- |
| **MI > 0.05** | 455 | 483 | 15 | 3% |
| **MI > 0.1** | 158 | 482 | 16 | 3% |
| **MI > 0.15** | 82 | 483 | 15 | 3% |
| **MI > 0.2** | 51 | 477 | 21 | 4% |
| **MI > 0.3** | 17 | 464 | 34 | 7% |

Fig. 3b shows the MI between the genes and the clusters and by setting a threshold in which genes with MI above it determines which genes are selected as input features for building the SVM (Fig. 3c). The gene selection step helps to remove many irrelevant genes which improves the classification accuracy. As can be seen in Table 4 setting higher threshold to the MI value leads to lower number of informative genes that are fed to the classifier and influence its performance. Using a MI threshold above 0.15, leads to 82 useful genes without damaging the classifier performance.

### Multiclass Support vector machine (SVM)

In machine learning SVM is a supervised learning model used either to classification or regression analysis, introduced in 1992 by Boser, Guyon and Vapnik (Boser et al., 1992). Given a labelled training data, an SVM classifies it by finding the best hyperplane that separate all the data points of one class from the other class. The best hyperplane for an SVM means the one with the largest margin between the two classes. The support vectors are the data points that are on the margins of the separating hyperplane. New data points are then mapped into the same space and predicted to belong to a specific class based on which side of the hyperplane they fall. It often happens that the sets to discriminate are not linearly separable in a finite dimensional space. In that case a kernel function is used to map the original finite dimensional space into a much higher dimensional space, making the separation easier in that space. The selection of an appropriate kernel function is important, since it defines the space in which the training set will be classified. Exploring of the different kernel function can be found in our manuscript in preparation, here we show the result of the linear kernel function.

The classification problem in this work is multiclass classification rather than a binary classification. To face this problem, the dominant approach is to reduce the single multiclass problem into multiple binary classification problems. Using the R package e1071 version 1.6.8 (Meyer et al., 2017), the “one-against one” approach was selected in which *n*(*n* − 1)/2 binary classifiers are trained where *n* is the number of classes (in this work *n* = 4); the appropriate class is assigned by the majority output of a voting scheme.

Choosing SVM in this work as a classifier was due to its high accuracy and its ability to deal with high dimensional data as was proven previously in large scale image classification and gene expression data (Abdullah et al., 2011; Jiang et al., 2007; Lin et al., 2011; Vanitha et al., 2015).

To train the SVM and test the its performance a leave one out cross validation (LOOCV) method was used. In this method, the train data is *N* − 1 cells, where *N* is the total number of cells in the embryo data (*498* cells) and the remaining *N*^*th*^ cell is used for testing the model and the same is repeated *N* times such that each cell is tested, classified and contributing to the model performance (Fig. 3c). The informative genes that passed the MI threshold were used as input features to the SVM. The LOOCV method makes the best use of the available data, especially when the number of samples is small (*498* cells), and avoids the problem of random selection (Ben-Dor et al., 2000).

### Predicting the class of the in vitro cells

1. Selecting the CLE cells in the same way it was done in the embryo data.
2. Selecting the same informative genes that were used to build the SVM on the embryo data.
3. Inserting the expression matrix of the in vitro cells as an input to the SVM.
4. The output is the probabilities of each cell to be assigned to any of the 4 trained clusters.
5. Choosing the dominant class that the cell was assigned in agreement with the maximum probability out of the 4 probabilities (see the plot under Step 6 in Fig. 4a).
6. Since the true classification of the cells is not known and since there might be some hidden classes in the in vitro data that were not trained using the embryo data, a harsh constrain needs to be taken: only the cells with minimum probability of 0.8 to be assigned to the dominant class are proceeded to the next step (see the probability plot under Step 6 in Fig. 4a: probability of 0.8 is indicated by the red line).
7. Classification results: only the qualified cells from the previous step are assigned to any of the 4 classes.

### Pseudotime analysis

The cells from the in vitro samples: ES-NMP, Epi-CE, Epi-CE-T and Epi-NMP, that were classified to the 4 classes (the qualified output cells from the SVM pipeline), went through a pseudotemporal cell ordering. For pseudotime reconstruction of single cell RNA-seq data there are not a lot of available tools that have been systematically tested and have easily accessible software. Moreover, in this work we are analysing a heterogeneous cell population of different conditions rather than cells from a time course experiment, hence the supervised pseudotime reconstruction approaches are not applicable and one should rely on unsupervised methods. We decided to use TSCAN, the Biocounductor R package version 1.16.0, (Ji and Ji, 2017), since it demonstrated reliable unsupervised pseudotime reconstruction results compared to alternative methods (Ji and Ji, 2017).

TSCAN first clusters the cells then it builds a minimum spanning tree to connect the clusters. The branch of this tree that connects the largest number of clusters is the main branch which is used to determine the psedotime order of the cells. This algorithm does not detect starting or ending points and prior biological information is needed to understand the start of the pseudotime order. The pseudotime order might represent the underlying developmental trajectory.

### Defining the highly expressed genes in the 2 pseudotime ranges of class 4

1. The cells in class 4 were split to 2 groups based on their pseudotime order: *t*_*p*_ ≤ 1250; 2500 ≤ *t*_*p*_ ≤ 3700.
2. Identifying the differential expressed genes between the 2 groups using the two sided Wilcoxon rank sum test. The P-value was corrected using the “BY” method of Benjamini and Yekutieli (2001). This method controls the false discovery rate and the proportion of false discoveries amongst the rejected hypotheses.
3. *4,569* differential expressed genes were detected by setting the adjusted *P* – *value* ≤ 0.01.
4. The mean expression of the *4,569* genes across the cells in each group was calculated.
5. Log2 fold between the mean expression of the 2 groups was calculated and the highly expressed genes in each group were defined as the genes that their log2 fold is above 1, resulting in *24* genes in the early pseudotime range and *178* genes in the later range (Table S5).
6. Using the ccRemover R package version 1.0.4 (Barron and Li, 2016) each gene from the identified highly expressed genes could be identified as cycling gene. *55%* of the highly expressed genes in the later pseudotime range group are defined as cycling genes, whereas the cells in the earlier range don’t show this enrichment (no cycling genes).

### Statistical test for controlling the sample size

**Table 5.** The number of the in vitro cells classified to the 4 clusters.

|  | Class 1 | Class 2 | Class 3 | Class 4 |
| --- | --- | --- | --- | --- |
| Number of cells | 1141 | 264 | 70 | 2036 |

Class 4 is approximately twice the size of class 1, and the node-like cells were assigned almost exclusivity to class 4. Hence, one might think that the different size of the classes might bias the finding of the node-like cells in class 4. The statistical test that was design in this case was aim to control for the size of the classes: randomly 570 cells (half of class1) were selected from class 1 and class 4, and the null hypothesis is that there is no difference in the number of the node-like cells between class 1 and class 4. This step was repeated 1,000 times, resulting that in 1,000 of the cases class 4 contained more node cells than class 1, means that the calculated *p* – *value* < 0.001 and the null hypothesis was rejected.

### Culturing Nodal-YFP cells and ubiquitous-tomato cells

Nodal::YFP and Sox17::GFP Ubiquitin::Tomato cells were cultured under the Epi-CE protocol, we name these cells Epi-CE Nodal and Epi-CE RFP (for red fluorescent protein) respectively. The Epi-CE RFP were further grown to make Epi-NMP (Epi-NMP RFP). After two days of culturing Epi-NMP RFP, we plated a mixture that consists of 50% Epi-NMP RFP positive RFP cells and 50% of positive YFP cells of Epi-CE Nodal, at a total dense of 5×10^4^ cells/cm^2^ (Fig. S8a-b). Since after sorting the cells they might be in stress, we decided to culture the mixture for 4 days and not for the normal period of 2 days to let the cells to recover. The mixture was grown in N2B27 supplemented with 20ng/ml FGF2 and 3μM Chiron to make Epi-meso2 (EM2), until sorting the cells to RFP positive (sample named: EM2-RFP+4d, contains only the Ubiquitin::Tomato cells) and RFP negative (sample named: EM2-RFP-4d, contains Nodal::YFP cells and might contain Ubiquitin::Tomato cells that didn’t express RFP, see Fig. S8c). This population of cells were compared to the EM2-RFP4d, which are 100% cells of Epi-NMP RFP plated in a flask and cultured for 4 days in N2B27 supplemented with 20ng/ml FGF2 and 3μM Chiron to make Epi-meso2. Total RNA was isolated from the 3 samples: EM2-RFP4d, EM2-RFP+4d and EM2-RFP-4d using Trizol. First strand cDNA synthesis was performed with Superscript III system (Invitrogen) and the quantification of double-stranded DNA obtained with specific genes designed primers, using QuantiFast SYBR Green PCR Master Mix (Qiagen) and the standard cycler program (Qiagen RotorGene Q). The qPCR was done in technical triplicates. The primers that have been used are available in Table 6. Expression values were normalized against the housekeeping gene *Ppia*. Here are the steps to calculate the normalized gene expression values:

1. Identifying the C_t_ (threshold cycle) for each gene (technical triplicates) and calculating the expression values (2^−Ct^).
2. Calculating the average and the standard deviation (std) for each gene from the triplicate expression values.
3. Dividing the average and the std of each gene in the expression value of *Ppia*.
4. The gene expression across the different conditions was scaled between 0 to 1.

**Table 6.**
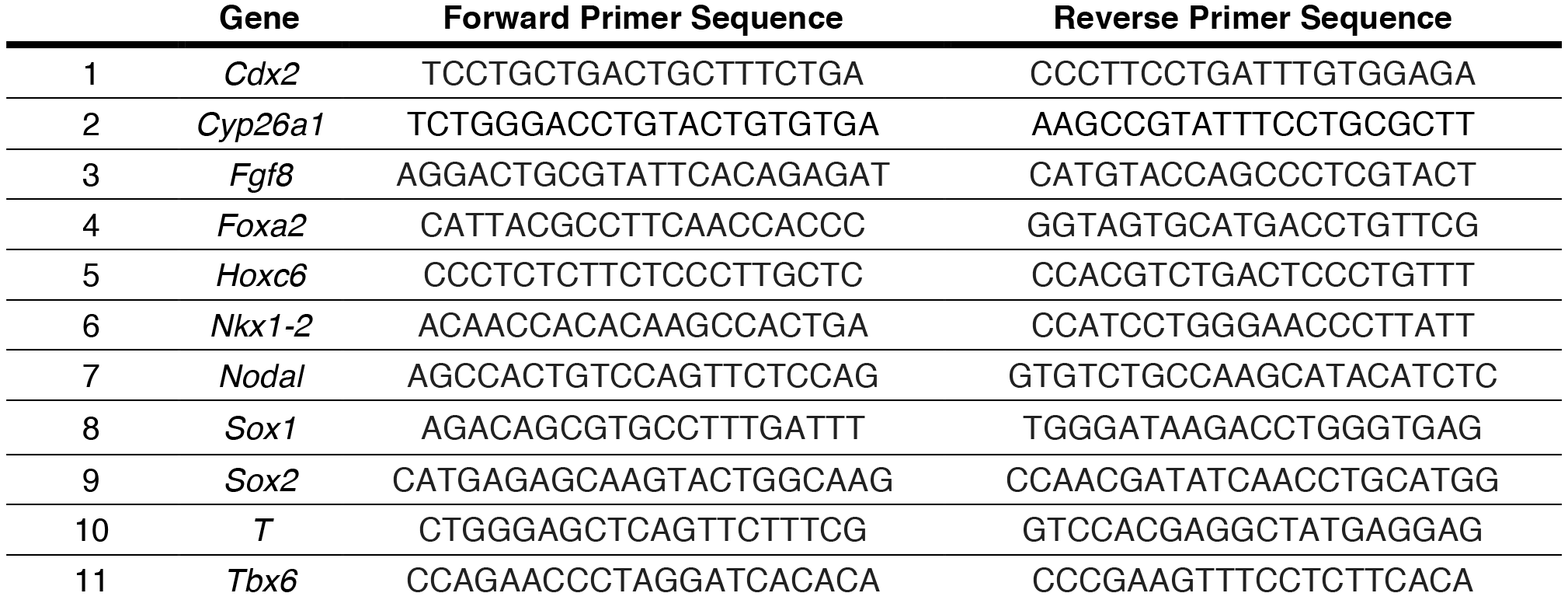
Primer sequences used for qRT-PCR.

### Cell sorting

Epi-CE Nodal cells were sorted according to their YFP positive fluorescence in a MoFlo sorter (Beckman Coulter) using 488nm laser with emission filter of 530/40 (Fig. S8a) and Epi-NMP RFP cells were sorted according to their RFP positive fluorescence using 647nm laser with emission filter of 610/20 (see Fig. S8b). Cells were collected, counted and replated in N2B27 supplemented with 20ng/ml FGF2 and 3μM Chiron medium to make the mixture of 50% Epi-CE Nodal YFP positive cells with 50% of Epi-NMP RFP positive cells, as it was described above. After 4 days, the mixture was sorted to RFP positive and negative cells in the MoFlo sorter using the same laser and filter sets mentioned above (Fig. S8c).

